# SeaB is a conserved *Salmonella enterica* extracellular matrix binding protein involved in biofilm formation and infection

**DOI:** 10.64898/2026.01.14.699485

**Authors:** Rochelle M. Da Costa, Jessica L. Rooke, Faye C. Morris, Zihao Yang, Zheng Jie Lian, Amanda E. Rossiter, Jeffrey A. Cole, Brian Forde, Adam F. Cunningham, Ian R. Henderson

## Abstract

*Salmonella enterica* is a leading cause of gastroenteritis worldwide. Exacerbating this issue is the emergence of multi-drug-resistant strains, posing a major threat to human health. Type 5 secretion system proteins play a major role in virulence and are viable vaccine targets. However, only a limited number of these proteins have been functionally characterized to date. In this study, we characterized SeaB, which belongs to the Type 5a secretion system. We demonstrated that SeaB is localized to the cell surface and is involved in binding to the extracellular matrix. Our results indicate that SeaB is involved in aggregation, biofilm formation and contributes to virulence. Furthermore, immunization with SeaB elicits antibodies and provides protection against *Salmonella* challenge in a mouse model of infection.

## INTRODUCTION

*Salmonella enterica* is a broad host range pathogen of animals and humans. Infection occurs through ingestion of contaminated food and water. Clinical disease is characterized by two major syndromes: systemic typhoid fever and enteritis. While *S. enterica* serovar Typhi is a human-restricted pathogen and is the predominant cause of typhoid fever, *S. enterica* Enteritidis and Typhimurium are non-typhoidal *Salmonella* (NTS) serovars and the main causative agents associated with enteritis in a wide variety of hosts [1]. In contrast to the self-limiting gastroenteritis in healthy individuals, invasive non-typhoidal *Salmonella* (iNTS) is one of the major causes of bacterial bloodstream infections in young children, elderly, immunocompromised and HIV-infected individuals in Sub-Saharan Africa [1, 2]. Strains associated with iNTS infections are primarily from serovars Typhimurium and Enteritidis [3]. In all *S. enterica* infections, key facets of pathogenesis are the ability to interact with host cells in the intestine and to develop an asymptomatic persistent infection in animals and humans, which provides a reservoir from which bacteria are shed in faeces to cause recurrent infections and outbreaks.

The high prevalence of antibiotic-resistant *S. enterica* infections has led the WHO to categorise this as a high priority pathogen for which new treatments and preventives are urgently required [4, 5]. With limited treatment options and the absence of new antibiotics in the development pipeline, vaccines that prevent disease onset or interrupt transmission by blocking persistent colonisation represent a compelling strategy for dealing with *Salmonella* infections [3]. Outer membrane proteins play a key role at the interface between the pathogen and host, serving as targets for antibodies and making them promising candidates for vaccine development. Gaining a deeper understanding of the surface-expressed virulence factors in *S.* Typhimurium is crucial for elucidating mechanisms of bacterial virulence and for identifying new potential vaccine targets.

Here we characterized SeaB (***S****almonella **e**nterica* **a**utotransporter involved in **b**iofilm formation), a surface protein conserved across all the different *Salmonella* serovars. SeaB is a member of the Type 5 secretion system (T5SS), which are found widely among Gram-negative bacteria. SeaB belongs specifically of the T5aSS subfamily [6–10]. Proteins belonging to this subclass are comprised of an N-terminal Sec-dependent signal sequence, a passenger domain and a highly conserved β-barrel domain [11–13]. The β-barrel is inserted into the outer membrane via the Bam complex with the aid of various periplasmic chaperones, and facilitates secretion of the passenger domain to the cell surface [14, 15]. With limited exceptions, the passenger domains of the T5aSS proteins adopt a β-helix conformation on the bacterial surface, the folding of which is facilitated by an autochaperone domain and the β-barrel [16]. The β-helix varies in length but is sufficiently long in each bacterium to display the functional component of the passenger domain beyond the LPS layer of the outer membrane [17].

T5SS proteins have been shown to be involved in adhesion, aggregation, biofilm formation and virulence [18]. Several are also components of licenced human vaccines [19]. Currently five T5SS proteins have been identified in *S.* Typhimurium; ApeE, MisL, ShdA and SeaB (previously YaiU), which are member of the T5aSS subfamily, and SadA which belongs to the T5cSS subfamily of trimeric autotransporters. ApeE is a lipase whose biological function is unknown [20]. MisL is encoded on *Salmonella* pathogenicity island 3 (SPI-3) and is involved in long term intestinal colonisation by binding to ECM fibronectin [21, 22]. ShdA is a large outer membrane protein that binds fibronectin and is involved in prolonged faecal shedding [23]. SadA is involved in biofilm formation, aggregation, adhesion and immunisation with this protein provides protection against *Salmonella* infection [24]. Importantly, multiple studies have shown that extracellular matrix (ECM) binding and biofilm formation are vital mechanisms of *Salmonella* persistence in mouse models of infection suggesting important roles for these proteins in establishing chronic infections across different species [9, 10]. However, the *Salmonella* T5aSS protein SeaB has not been characterized previously. A study by Chaudhuri *et al.* using the transposon directed insertion site sequencing (TraDIS) observed that mutations in *seaB* resulted in attenuated virulence in chickens, pigs and cattle [25]. Other studies have shown that EhaB, a SeaB homologue in enterohemorrhagic *Escherichia coli,* is involved in biofilm formation and adhesion to ECM components, indicating a similar role for SeaB in *Salmonella* [26]. Thus, here we analysed the conservation of SeaB in the *Enterobacteriaceae*, and characterized its role in adhesion, aggregation, biofilm formation and virulence. We also assessed its potential as a vaccine candidate.

## MATERIALS AND METHODS

### Bioinformatic analyses

The amino acid sequence of SeaB (SL1344_RS01910) from *S*. Typhimurium strain SL1344 was used as a query. SeaB homologues in other bacterial species were identified by BlastP searches using the RefSeq database on NCBI [27]. Homologues were identified as having an e value of <0.001 with at least 30% sequence identity. Phylogenetic trees were drawn using IQ-TREE v2.3.6 [28] on Galaxy [29], with maximum likelihood and >1000 bootstraps. Trees were annotated and edited using iTol v6.9.1 [30]. The genetic loci of SeaB homologues in related species were downloaded from BioCyc [31–33] and edited using Adobe illustrator.

### Bacterial strains, plasmids, and growth conditions

Strains and plasmids used in this study are listed in Table 1. Bacteria were cultured in Luria Bertani broth (LB) at 37°C with shaking unless stated otherwise. If required, the culture medium was supplemented with 100 µg/ml carbenicillin, 50 µg/ml kanamycin or 0.5 mM isopropyl-β-D-thiogalactopyranoside (IPTG).

**Table 1:** List of strains and plasmids.

| Name | Description | Reference |
| --- | --- | --- |
| **Strains** |  |  |
| *S. Typhimurium* SL1344 | *Salmonella enterica* serovar Typhimurium | [61] |
| SL1344 *seaB*::aph | *SeaB* gene replaced with kanamycin resistance cassette | This study |
| *S. Typhimurium* SL3261 | Δ*aroA* derivative of SL1344 | [62] |
| SL3261 *seaB*::aph | *SeaB* gene replaced with kanamycin resistance cassette | This study |
| *E. coli* DH5α | High efficiency competent *E. coli* derived from DH5α | New England Biolabs |
| *E. coli* BL21 DE3 | T7 expression *E. coli* strain for protein expression | New England Biolabs |
| **Plasmids** |  |  |
| pET22b+ | T7 expression vector | Novagen |
| pREP4 | Repressor plasmid for controlled expression of pQE vectors | Qiagen |
| pQE60 (modified) | Modified expression plasmid | [24] |
| pET22b - OmpA | OmpA signal sequence (ss) cloned into pET22b+ between *NcoI* and *NdeI* restriction sites | This study |
| pET22b-OmpAss-SeaB | *SeaB* cloned with N-terminal His tag between *NotI* and *KpnI* sites of pET22b+ | This study |
| pQE60 SeaB | *SeaB* gene cloned between *NdeI* and *HindIII* sites | This study |

### Molecular biology techniques

Genomic DNA and plasmid DNA were extracted using the Qiagen gDNA kit and Qiagen mini prep kit as per manufacturer’s instructions. Samples were quantified by nanodrop. Primers used in this study are listed in Table S1. Fragments used for cloning were amplified using Phusion High-Fidelity DNA Polymerase (New England Biolabs) or MyTaq (Bioline) as per manufacturer’s instructions. PCR products were purified using the Qiagen PCR purification kit as per manufacturer’s instructions unless stated otherwise. FastDigest enzymes (Thermo Scientific) and T4 ligase (NEB) were used for cloning reactions and constructs were transformed into NEB 5-alpha Competent *E. coli* (High efficiency) before plating on LB agar with appropriate antibiotics. The *seaB* mutant strains were generated as described previously [34].

### Protein expression and whole cell protein extraction

For protein expression, *seaB* was cloned into pET22b+ plasmid. The resulting plasmid was transformed into BL21 DE3 or BL21 (DE3) P2. Transformants were incubated overnight at 37°C with 100 µg/ml carbenicillin. The following day, 4 L of 2 x LB was inoculated with a 1:100 dilution of overnight cultures and grown at 37°C with aeration. Cultures were grown to an OD_600_ of ∼0.6 and induced with 50 µM isopropyl β-D-1-thiogalactpyranoside (IPTG). Induced cultures were grown for 16 h with aeration at 16°C. Whole cell protein fractions were prepared from 10^9^ CFU from overnight cultures. Bacteria were collected by centrifugation at 13,000 *g* for five min and re-suspended in 2 x Laemmli sample buffer (Sigma Aldrich). This mixture was boiled for five min at 100°C and centrifuged at 13,000 *g* for two min prior to loading onto SDS-PAGE gels.

For the purification of membrane localised SeaB, 3 L of induced cultures were pelleted by centrifugation at 5,000 *g* for 10 min at 4°C. The pellet was then resuspended in ice-cold 20 mM sodium phosphate (pH 7.4), 500 mM NaCl and 2 mM PMSF. Cells were lysed using the Avestin C3 and unbroken cells and debris were collected by centrifugation at 5,000 *g* for 10 min at 4°C. The membrane was isolated from the soluble lysate fraction by centrifugation at 50,000 *g* for 45 min at 4°C. The membrane pellet was then resuspended in 20 mM sodium phosphate pH 7.4, 500 mM NaCl, 2% n-Dodecyl-β-D-maltoside (DDM) detergent and 20 mM imidazole (1 ml of buffer per 100 mg of the pellet). The membrane pellet was left to solubilise overnight at 4°C on end-over-end tube rotator. The following day, the insoluble fraction was pelleted by centrifugation at 50,000 *g* for 60 min at 4°C. The soluble fraction was coated in Ni-NTA agarose beads (Invitrogen) for one hour at 4°C on an end-to-end rotator and applied to an Econo-Pac chromatography column (Bio-Rad). The columns were washed with 100-300 ml of wash buffer (50 mM sodium phosphate pH 7.4, 10 mM imidazole, 400 mM NaCl and 0.03% DDM) before being eluted in 10 ml of elution buffer (50 mM sodium phosphate pH 7.4, 500 mM imidazole, 500 mM NaCl and 0.03% DDM). Eluted fractions were analysed by SDS-PAGE and fractions containing the purified full-length protein was buffer exchanged through snakeskin dialysis tubing (Thermo Fischer Scientific), with a molecular weight cut-off of 3.5 kDa, into a buffer containing 10 mM sodium phosphate pH 7.4, 250 mM NaCl. The purified proteins were stored at 4°C until further use. Concentration of the purified protein was measured using the Pierce BCA protein assay kit (Thermo Fischer Scientific) and the endotoxin levels in the purified protein preparations were tested using the Pierce Endotoxin Quant kit (Thermo Fischer Scientific). To assess the folded state of the proteins heat modifiability assays were performed. Briefly, 10 µl of purified protein was added to 1.5 ml Eppendorf tubes and 90 µl of 2 x Laemmli sample buffer was added to each tube. One sample was heated for 15 min at 100°C while the other was incubated at room temperature prior to SDS-PAGE analysis. Localization of SeaB to the cell surface by immunofluorescence was performed as described previously [24, 26]. Purified protein was used for immunization studies in mice, and polyclonal antibodies against SeaB were raised in rabbits following immunization with purified protein.

### Functional assays

To test binding to ECM components, 96-well NUNC microwell plates were coated with 100 µl of 2 µg/ml of collagen, 2 µg/ml fibronectin and 2 µg/ml of bovine serum albumin (BSA) as a control and left at 4°C overnight or for one hour at 37 °C. The following day, plates were washed three times with 1 x PBS containing 0.05% TWEEN 20 (v/v). Next, the plates were coated with 100 µl of 1 µg/ml of purified protein. Following incubation with the purified proteins, unbound proteins were washed and 1:2000 dilution of anti-His primary antibody (Jomar Life Research) was added to the wells. The plates were incubated at 37°C for 1 hour and washed again before the addition of 1:2000 IgG conjugated with alkaline phosphatase (Merck Life Science). To measure the colorimetric indirect ELISA response, SIGMAFAST *p*-nitrophenyl phosphate (Sigma Aldrich) was prepared as per manufacturer’s instructions and 100 µl of substrate was added per well. The plate was incubated at room temperature for 30 min and the absorbance OD_405nm_ was measured using a TECAN plate reader (Infinite M Plex). Autoaggregation and biofilm assays were performed as described previously [35]. Cell culture and adhesion/invasion experiments were performed as described previously in [35, 36].

### Murine infection studies

Mice used in this study were approved by the animal ethics committee at the University of Queensland under approval number IMB/382/19 and 2021/AE001097. All experiments used C57BL/6J (non-genetically modified) male mice, 6-8 weeks of age, purchased from the Animal Resource Centre (ARC)/Ozgene located in Perth, Australia. C57BL/6 male mice (6-8 weeks) were infected via the intraperitoneal route with 1000 CFU/dose *S.* Typhimurium SL1344 or 5 x 10^5^ CFU/dose SL3261; and 10^8^ CFU/dose *S.* Typhimurium SL1344 via the oral route. Mice were culled by carbon dioxide asphyxiation at day 5 post challenge with SL1344; days 7, 21 and 35 post challenge with SL3261 and day 2 post challenge with SL1344 for immunization and challenge studies. Livers, spleens, and gall bladders were collected from mice at the time of euthanasia. Organs were homogenised using the Bullet blender storm pro homogenizer (Next Advance) and subsequently serially diluted and plated on LB agar. After overnight incubation the CFU/organ were enumerated.

To study the protective effect of immunizing with SeaB, 100 µg of the purified protein was prepared in sterile PBS . Imject Alum (Thermo scientific) was added dropwise to the protein preparation so that the final volume ratio of alum to protein was 1:3 and mixed using end-to-end tube rotator for 30 min at room temperature. Mice were immunized intraperitoneally with 10 µg of the purified protein on day 0 and day 30. Mice were anaesthetised with isoflurane for blood collection via tail bleed (typically 50-100 µl) at days 0, 8, 28 and 43. Blood was incubated at 37°C for 1 h and centrifuged at 1,000 *g* for 15 min at 4°C. Organs were harvested and bacterial numbers enumerated as above.

### Enzyme linked Immunosorbent assays

Blood was collected from mice via tail bleeds or cardiac puncture and clotted at 37°C for 1 h. Blood was centrifuged at 1000 *g* for 15 min at 4°C and sera were transferred to fresh eppendorfs and were stored at -80 until further use. NUNC 96 plates were coated with 1 µg of purified protein prepared in coating buffer (0.015 M sodium carbonate, 0.035 M sodium bicarbonate pH 9.6). Plates were incubated at 37°C for 1 h or overnight at 4°C.The following day plates were washed with wash buffer (3x) (0.1 M PBS and 0.05% Tween 20) and 200 µl of blocking buffer (0.1 M PBS and 1% BSA) added to each well for 1-2 h at 37°C. Plates were washed with wash buffer (3x) and the required volume of buffer and test serum were added using a starting dilution of 1:20 and diluting 3-fold. The plates were incubated at 37°C for 1 h. Plates were washed again in wash buffer (3x) prior to the addition of secondary antibody, anti-mouse IgG AP (1:2000) or anti mouse IgM AP (1:2000) and incubated at 37°C for 1 hour. SIGMAFAST p-Nitrophenyl phosphate substrate (Tris buffer tablet and PNPP tablet) was prepared in distilled water in accordance with manufactures instructions, before being added to each well and incubated at room temperature. After incubation, the OD_405nm_ was measured using a TECAN plate reader at regular intervals. The antibody titres were determined by generating an ELISA curve of the dilution of the sera that were parallel and plotted as relative antibody titres.

For whole cell ELISAs, bacteria were grown to OD_600nm_ of 0.6. Cells were centrifuged at 4,500 *g* for 10 min and washed three times in coating buffer (0.015 M sodium carbonate, 0.035 M sodium bicarbonate pH 9.6). The cell pellet was resuspended to a final volume of 10 ml in coating buffer. NUNC 96-well plates were coated with 100 µl of the cell suspension and incubated overnight at 4°C. The following day plates were washed with wash buffer (0.1 M PBS and 0.05% Tween 20). The rest of the protocol was as per the ELISA protocol as described above.

### Figures and statistics

Figures were made using Adobe Ilustrator. GraphPad prism (version 9.3.1) was used for all the statistical analysis. Mann Whitney non-parametric U test, two-way ANOVA corrected for multiple comparisons were used. The details of the statistical analysis are stated in the figure legends.

## RESULTS

### SeaB is conserved across Enterobacteriaceae and Yersiniaceae

Genomic analysis of *S. enterica* serovar Typhimurium strain SL1344 identified a 2,934-bp open reading frame, designated *STM0373*, predicted to encode a 978-amino acid autotransporter protein of the T5aSS subfamily, herein referred to as SeaB. *In silico* analysis revealed a canonical Sec-dependent signal peptide, with a predicted cleavage site following residue 27. The passenger domain (residues 28–728) comprises a 701-amino acid region containing a Pertactin-like motif predicted to adopt a single-stranded, right-handed β-helix structure, along with a C-terminal autochaperone (AC) domain. The translocator domain (residues 729–978), indicated by Pfam (PF03797) as an autotransporter β-domain, is predicted to form a 12-stranded outer membrane embedded β-barrel with an α-helical linker. Structural modelling of SeaB suggests surface exposure of the AC domain and luminal positioning of the α-helix within the β-barrel, consistent with the architecture of characterized T5aSS autotransporters (Fig. S1). In contrast to the solved structures of Pertactin, Antigen 43 and other T5aSS proteins, the passenger domain of SeaB is predicted to be decorated along one axis of the β-helix with disordered loops.

To identify orthologues of SeaB, the full-length amino acid sequence was queried against the RefSeq database of bacterial genomes using NCBI BLASTP. Candidate sequences were subsequently filtered based on domain architecture congruent with that of a T5aSS protein and a minimum of 30% sequence similarity. Multiple sequence alignment of SeaB homologues was performed, and a maximum-likelihood phylogenetic tree was constructed. These analyses revealed that SeaB orthologues are broadly found within the *Enterobacteriaceae*, including *E. coli* and *Citrobacter rodentium*, but are notably absent in *Klebsiella pneumoniae* (Fig. 1A). In all cases, the *seaB* gene was located adjacent to the *hemB* locus and upstream of *iprA* (Fig. 1B).

**Figure 1.**
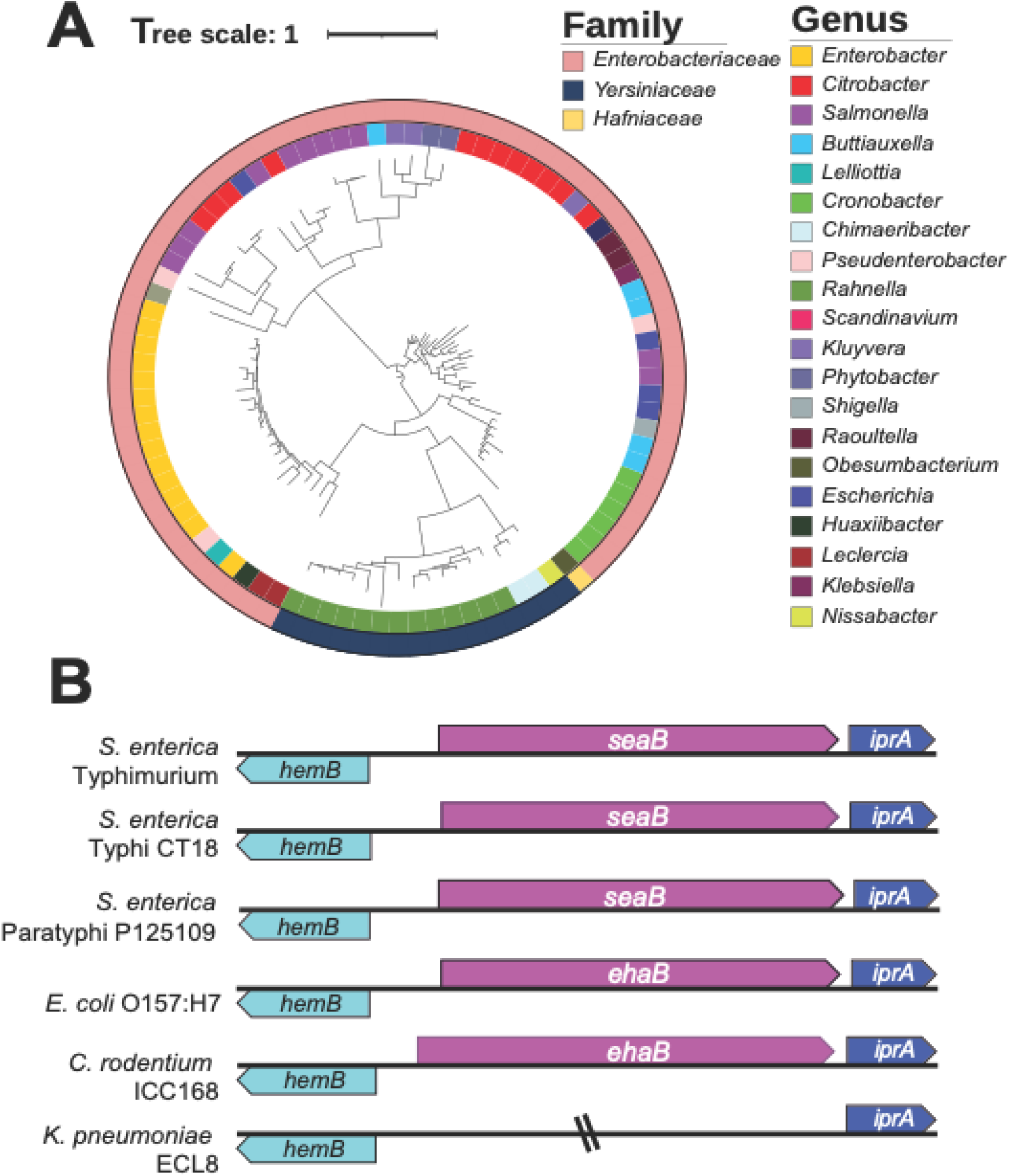
SeaB conservation. (A) Phylogenetic tree SeaB homologues retrieved from BlastP searches using the refseq database. The tree was generated using IQTree with >1000 bootstraps and edited in iTol. The inner ring denotes genus and the outer denotes Family. (B) Neighbourhood analysis of SeaB homologues from closely related Enterobacteriaceae.

### SeaB is surface localized

T5aSS proteins are found in Gram-negative bacteria and can function as virulence factors. These virulence factors are surface exposed, form an interface between the pathogen and the host, and are antibody targets [19]. SeaB shares key structural features with other T5aSS autotransporters, including a predicted C-terminal β-barrel domain and surface exposed N-terminal passenger domain.

To assess the subcellular localization of SeaB, the protein was heterologously expressed in *E. coli* BL21(DE3) using the pET22b expression system (Fig. S2A - D). The presence of SeaB in the outer membrane fractions was confirmed by Western Blotting using anti-SeaB antibodies, where a band (107 kDa) was observed only in fractions harbouring a plasmid expressing *seaB* (Fig S2E). No band was observed in the wild-type or *seaB* mutant strains. The observed band has a mass consistent with the predicted molecular weight of SeaB (103 kDa) after the signal sequence has been processed. Surface exposure of SeaB was evaluated by whole-cell ELISA and immunofluorescence microscopy.

*E. coli* BL21 strains harbouring either the empty vector (pET22b EV) or the SeaB expression construct (pET22b SeaB) were immobilized on ELISA plates and probed with anti-SeaB or anti-RNA polymerase (RNAP) antibodies. A significantly higher signal was observed in the SeaB-expressing strain when probed with anti-SeaB antibodies compared to the vector control (Fig. S2F), indicating surface localization of the recombinant protein. No significant signal was detected with anti-RNAP, confirming that only surface-exposed antigens were detected under these assay conditions. Surface localization of SeaB was further validated by immunofluorescence microscopy. The *S. enterica* serovar Typhimurium SL1344 wild-type strain showed low fluorescence, consistent with the inability to detect the protein in outer membrane fractions of the wild-type bacterium, and findings from the Hinton lab that indicated *seaB* was poorly expressed under standard laboratory conditions [37]. No fluorescence was detected in the *seaB* mutant strain. However, complementation of the mutant generated a strong fluorescence signal on the cell surface (Fig. S2G). These findings support the surface exposure of SeaB and indicate that SeaB is not processed after outer membrane localisation, consistent with its predicted function as a T5aSS autotransporter and potential virulence factor.

### SeaB mediates aggregation and biofilm formation

Many of the characterized T5aSS have been implicated in bacterial aggregation and biofilm formation [38, 39]. Additionally, EhaB, a SeaB homologue in *E. coli*, has been shown to contribute to biofilm development [26]. To assess whether SeaB mediates similar phenotypes, we investigated autoaggregation and biofilm formation in *S. enterica* serovar Typhimurium SL1344 wild-type (WT), a SeaB knockout strain (SL1344 *seaB::aph*), and a complemented strain (SL1344 *seaB::aph* carrying pREP4 pQE60-SeaB). *E. coli* BL21 strains carrying either an empty vector or overexpressing SeaB were included for comparison. No differences in autoaggregation were observed among the *Salmonella* strains (Fig. 2A). In contrast, *E. coli* BL21 overexpressing SeaB exhibited a pronounced autoaggregation phenotype relative to the vector control (Fig. 2B). In biofilm assays, none of the *Salmonella* strains, nor the *E. coli* BL21 empty vector control, showed detectable biofilm formation under the tested conditions (Fig. 2C). However, robust biofilm formation was observed in *E. coli* BL21 overexpressing SeaB following 16 hours of incubation at 37°C under shaking conditions (Fig. 2D).

**Figure 2.**
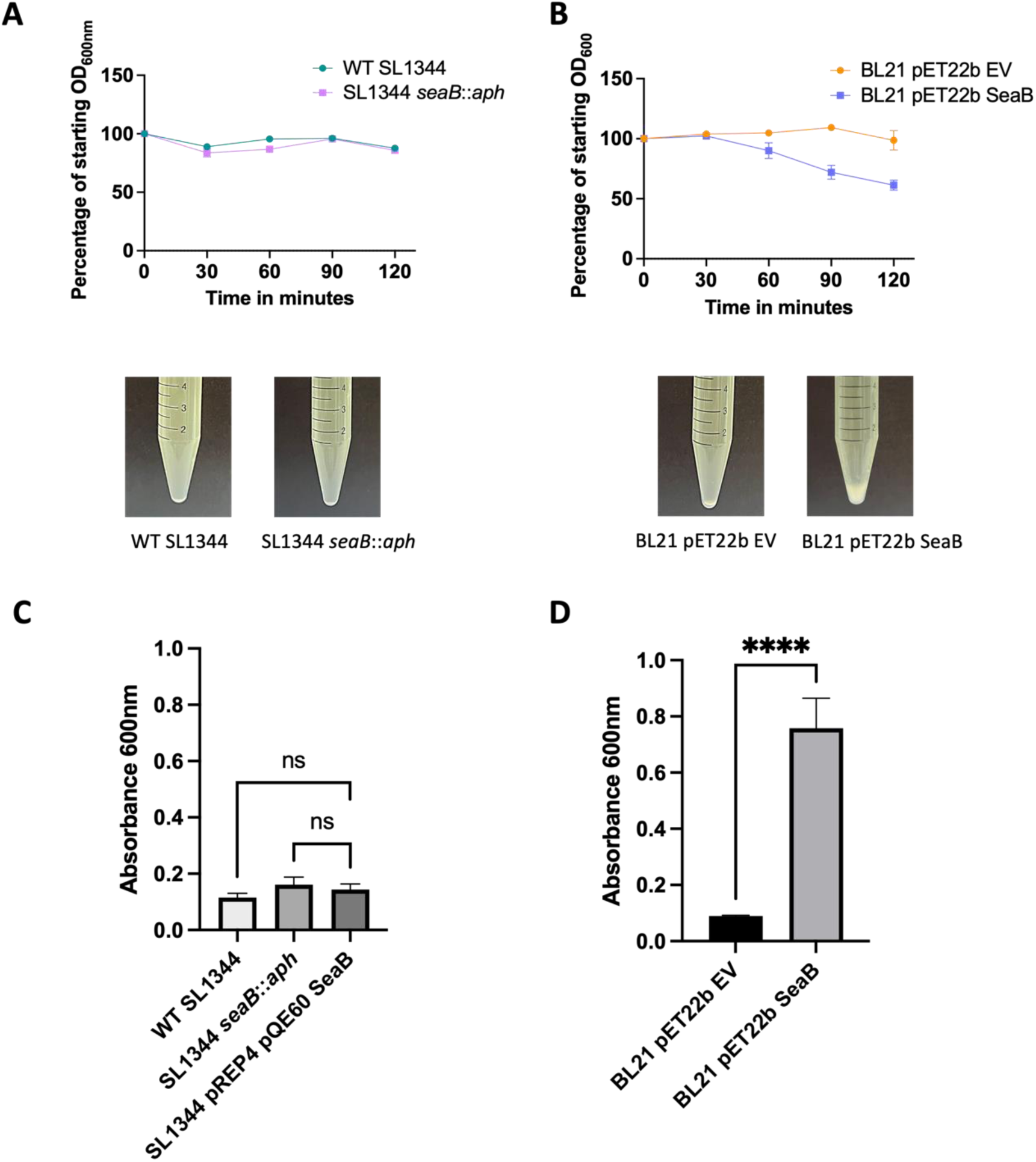
Autoaggregation and biofilm formation. Aggregation of bacterial strains (A) WT SL1344, SL1344 *seaB*::*aph*, (B) *E. coli* BL21 EV and *E. coli* BL21 pET22B SeaB. Images below show settling of the four strains under static conditions. Biofilm profiles of (C) *Salmonella* SL1344 strains (WT, *seaB*::*aph*, and pREP4 pQE60 SeaB, and (D) *E.coli* BL21 strains (EV and pET22B SeaB) after 16 hours of growth. Data (mean +SEM, n=3) from three independent experiments. Statistical significance was determined using a two-way ANOVA Tukey’s multiple comparisons test (ns p>0.05, and **** p<0.0001).

These findings indicate that SeaB can mediate aggregation and biofilm formation in a heterologous *E. coli* background, although this phenotype is not observed in *Salmonella* under the conditions tested.

### SeaB promotes binding to extracellular matrix components

In addition to their role in aggregation, several T5aSS autotransporters, including MisL and ShdA, have been shown to mediate binding to extracellular matrix (ECM) proteins such as collagen IV and fibronectin [22, 40], contributing to bacterial adherence and invasion of epithelial cells [38, 39]. To investigate whether SeaB facilitates similar interactions, we first assessed its ability to mediate adhesion to ECM components. As autoaggregation and biofilm formation phenotypes were obscured in *Salmonella*, subsequent functional assays focused on the *E. coli* BL21 strain overexpressing SeaB, where the phenotypes were more readily detectable. To evaluate ECM binding, *E. coli* BL21 expressing SeaB was incubated on plates coated with fibronectin, collagen III, or collagen IV. A significant increase in adherence was observed on all ECM-coated surfaces compared to the empty vector control (Fig. 3A). Among the substrates tested, the strongest binding was observed on collagen III, followed by fibronectin and collagen IV. These results confirm that SeaB mediates specific interactions with ECM components, supporting its proposed role as an ECM-binding adhesin.

**Figure 3.**
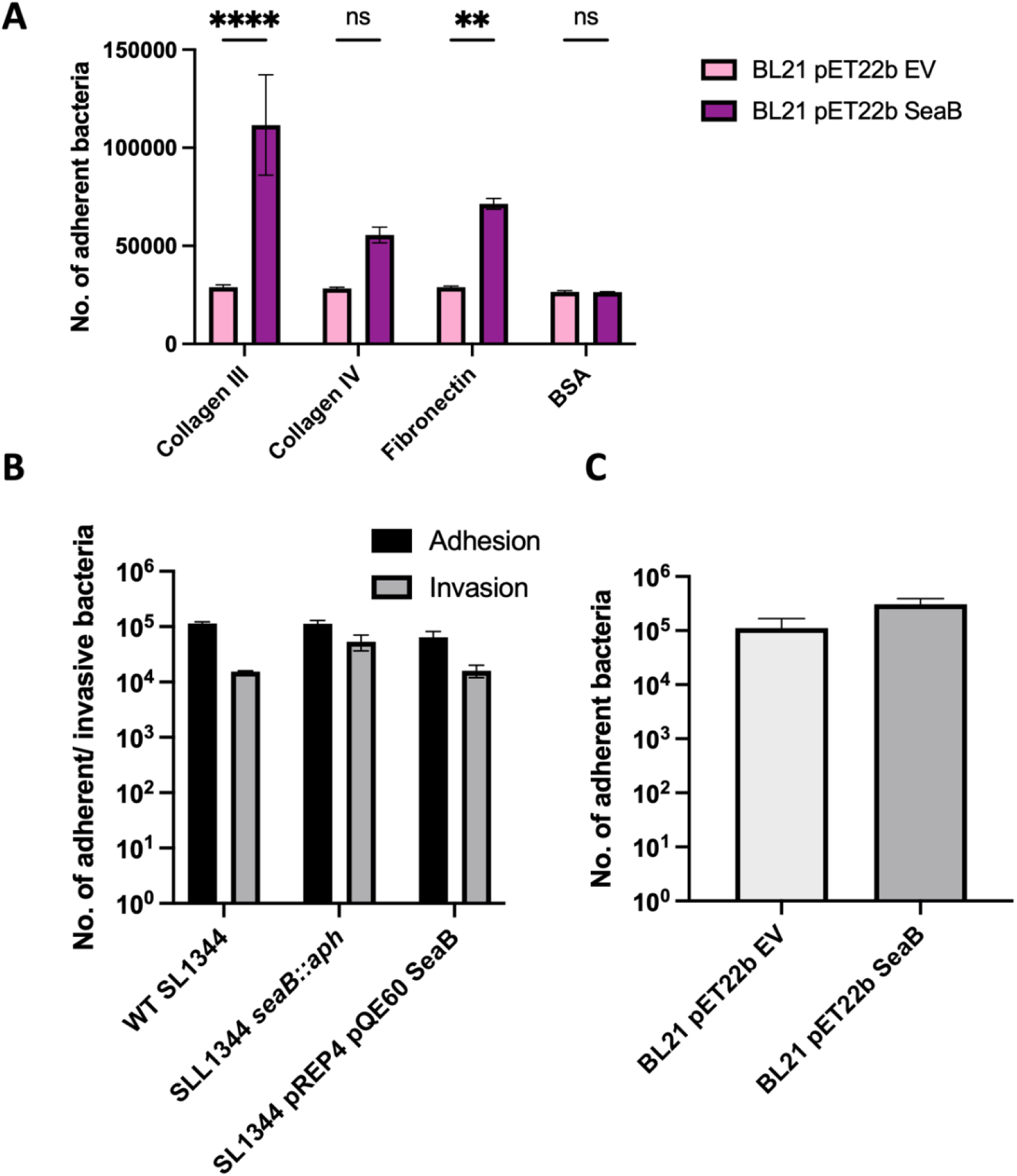
Adherence and invasion of SeaB. (A) ECM binding assay of *E. coli* BL21 EV and *E. coli* BL21 pET22B SeaB to collagen III, collagen IV, Fibronectin and BSA (control). Adherence and invasion of (B) *Salmonella* SL1344 strains (WT, *seaB*::*aph*, and pREP4 pQE60 SeaB, and (C) *E. coli* BL21 strains (EV and pET22B SeaB) to intestinal Caco-2 cells. Data (mean +SEM, n=3) are from three independent experiments. Statistical significance was determined using a two-way ANOVA multiple comparisons test (ns p<0.05, **p<0.01 and **** p<0.0001).

Subsequently, to determine whether SeaB contributes to host cell adhesion and invasion, we evaluated the interaction of *S. enterica* serovar Typhimurium SL1344 wild-type, a *seaB* deletion mutant, and a complemented strain with CaCo-2 epithelial cells. Adhesion and invasion assays revealed that deletion of *seaB* did not significantly alter either bacterial adherence or invasion relative to the wild-type strain. Furthermore, constitutive expression of SeaB from the pQE60 plasmid had no impact on this phenotype (Fig. 3B). Additionally, *E. coli* expressing SeaB did not exhibit enhanced adhesion compared to controls (Fig. 3C). Together, these results indicate that SeaB facilitates bacterial adherence to ECM proteins but may not play a direct role in adhesion to or invasion of epithelial cells.

### SeaB plays a role in virulence

Studies have shown that T5SS protein such as MisL and ShdA are involved in intestinal colonisation and persistent infection when mice are infected orally [22, 40]. To investigate whether SeaB has a role in infection, C57BL/6 mice were infected orally with 10^8^ CFU/dose of *S. enterica* SL1344 or the isogenic *seaB::aph* mutant. Bacterial colonisation of the liver, spleen, gall bladder and blood were determined on day 5 after challenge. A significant reduction in bacterial numbers was observed in the liver, spleen and blood at day 5 (Figure 4 A-E) post-infection with *seaB::aph* when compared to the wild-type strain. In contrast, there was no significant reduction in bacterial numbers in the gall bladder and ileum on day 5 post infection.

**Figure 4.**
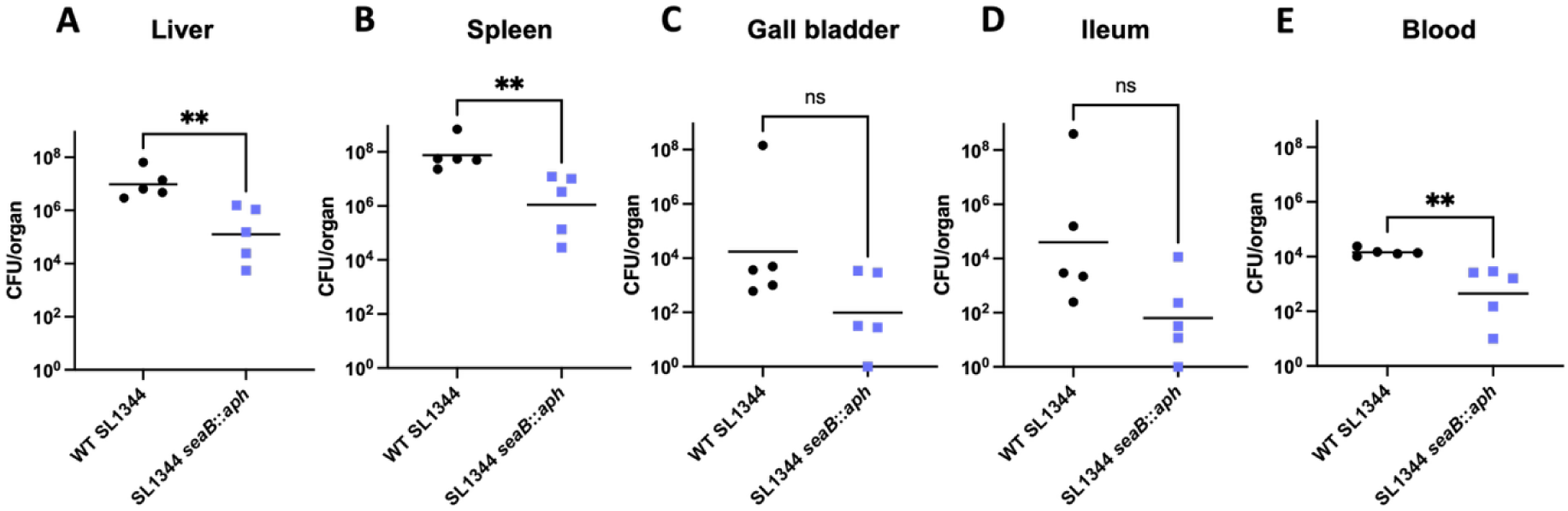
Infection dynamics after oral infection with *Salmonella*. Bacterial burdens in the (A) liver, (B) spleen (C) gall bladder and (D) blood of mice infected for 5 days with WT SL1344 (black closed circles) and SL1344 *seaB::aph* (purple closed boxes). Mice were culled at day 5 post infection. Statistical significance was determined using Mann-Whitney non-parametric test with correction for multiple comparisons (ns p>0.05, and ** p<0.01).

We next sought to determine whether a strain lacking SeaB was required for colonisation and persistence during late stages of systemic infection. To test this, C57BL/6 mice, which normally succumb to infection with wild-type *S.* Typhimurium, were infected intraperitoneally with an attenuated strain of *Salmonella* (WT SL3261) and the mutant strain (SL3261 *seaB::*aph) [41] and culled on days 7, 21 and 35 to determine bacterial burdens in the liver, spleen, and gall bladder. In contrast to oral infection, when mice were infected intraperitoneally in a chronic carriage model of *Salmonella* infection, a slight reduction in bacterial numbers post-infection with *seaB::aph* was seen at day 7 in the liver and days 7 and 35 in the spleen (Figure S3 A-C). However, no significant reduction in bacterial numbers was seen in the gall bladder. These results suggest that SeaB is required for oral infection of *Salmonella* but not chronic carriage.

### Immunization with SeaB protects against *Salmonella* infection

Given that SeaB is surface-exposed and appears to contribute to virulence-associated phenotypes, we next sought to determine whether immunization with purified SeaB could elicit a protective immune response in a murine model. To ensure that any observed protective effects were attributable solely to SeaB and not to contaminants from *Salmonella,* the protein was heterologously expressed and purified from *E. coli*. As protective antibody responses are typically directed against conformational epitopes, it was essential to confirm that the purified SeaB protein retained its native folded state prior to immunization. To assess this, we performed a heat modifiability assay, exploiting the known property of β-barrel domains in Type 5 Secretion System (T5SS) proteins, which unfold upon boiling and consequently exhibit altered migration on SDS-PAGE [42]. Boiled SeaB samples migrated further than their non-boiled counterparts (Fig. S2D), consistent with denaturation of a folded β-barrel structure. These results indicate that the purified SeaB protein retained its native conformation, supporting its suitability for use in immunization studies.

To determine whether SeaB can induce a protective immune response, mice were immunized intraperitoneally with 10 µg of purified SeaB adsorbed to alum on days 0 and 30. Control animals received PBS alone. A schematic overview of the immunization and challenge timeline is shown in Fig. 5A. Serum was collected on days 8, 28, and 43 to evaluate the development of SeaB-specific antibodies. Elevated levels of SeaB-specific IgM and IgG in immunized mice compared to PBS-treated controls were detected by ELISA on days 8 and 28, respectively (Fig. 5B), indicating successful induction of a humoral immune response. To assess the protective potential of this response, immunized and control mice were challenged intraperitoneally with *S. enterica* SL1344 on day 44.

**Figure 5.**
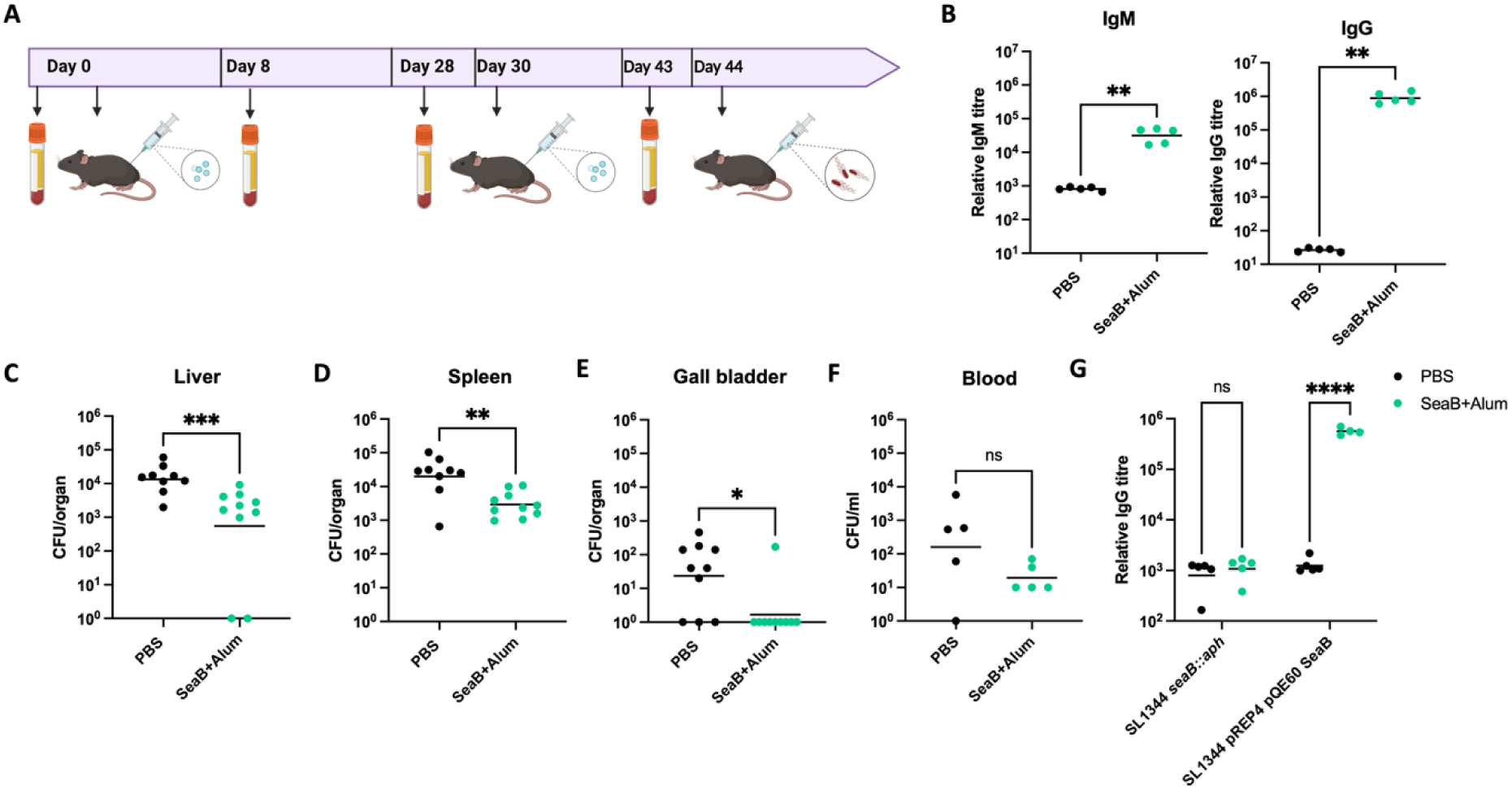
Immunization with purified SeaB. (A) Schematic overview of immunization experiment. 8-week-old C57BL/6 male mice were immunized with 10 µg of purified SeaB+Alum. Mice were challenged with WT SL1344 and culled on day 2 post infection. (B) IgM and IgG antibody titres determined by ELISA at days 8 and 43 respectively. (C) Bacterial burdens in the liver (D) spleen (E) gall bladder (F) blood from mice immunized with SeaB+Alum (green closed circles) and PBS controls (black closed circles). (G) Surface localisation of SeaB as measured by whole cell ELISA. Statistical significance was determined using a Mann-Whitney non-parametric U test and two-way ANOVA with Sidak’s multiple comparisons test (ns p>0.05, *p<0.05, ** p<0.01, *** p<0.001, ****p<0.0001).

Bacterial loads in the liver, spleen, and gall bladder were quantified two days post-infection. SeaB-immunized mice exhibited significantly reduced bacterial burdens in all three organs compared to control animals (Fig. 5C-F). These results demonstrate that immunization with purified SeaB elicits a specific antibody response and confers partial protection against systemic *Salmonella* infection.

To be protective, antibodies must recognize their antigen in a conformationally native state on the microbial surface [43]. To assess whether antibodies generated against purified SeaB could access the native protein on the bacterial surface, By coating ELISA plates with either a SeaB-deficient strain (*seaB::aph*) or the SeaB-complemented strain (SL1344 pREP4 pQE60-SeaB) minimal cross-reactivity with other *Salmonella* surface antigens was confirmed by the absence of significant IgG binding in the plates were coated with the SeaB mutant strain. In contrast, sera from immunized mice exhibited significantly higher IgG titres against the SeaB-complemented strain compared to sera from non-immunized controls (Fig. 5G). These findings indicate that immunization with purified SeaB induces antibodies capable of recognizing the native, surface-exposed form of SeaB on *Salmonella*, supporting its accessibility and immunogenicity *in vivo*.

## DISCUSSION

The T5SS is the most prevalent secretion system among Gram-negative bacteria and plays a pivotal role in bacterial pathogenesis. Proteins secreted via the T5SS contribute to key virulence-related processes including adhesion, autoaggregation and biofilm formation, making them attractive candidates for vaccine development [18]. Notably, some T5SS proteins are already incorporated in licensed human vaccines [44–46]. Despite their abundance and functional relevance, relatively few T5SS proteins have been experimentally characterized to date, and further research is needed to elucidate their roles in virulence and their potential as protective antigens.

In *S. enterica* serovar Typhimurium, only four T5SS proteins have been characterized to date: MisL and ShdA, both classified as Type 5a secreted proteins (T5aSS); SadA, belonging to the T5cSS subclass; and SinH, a member of the recently proposed T5eSS subclass [10, 21]. MisL and ShdA facilitate intestinal colonization, faecal shedding and long-term persistence in murine models through interactions with extracellular matrix (ECM) components [22, 39, 40]. In addition, MisL contributes to bacterial aggregation and biofilm formation, and immunization with MisL has been shown to reduce bacterial burdens following *Salmonella* challenge [38, 47]. Similarly, our previous work demonstrated that SadA promotes aggregation, biofilm formation and adhesion and can elicit protective immunity when used as a vaccine antigen [36].

Through comparative genomic analyses, we identified *seaB*, a previously uncharacterized gene in *S.* Typhimurium encoding a predicted T5aSS autotransporter. Orthologues of *seaB* are variably annotated in public databases (e.g., *yaiU* or *yaiT*) based on a corresponding pseudogene in *E. coli* K-12. Homologues of s*eaB* are widely distributed among closely related species such as *E. coli* and *C. rodentium*. The consistent genomic positioning of *seaB* across different taxa suggests that the gene originated prior to the divergence of *Salmonella*, *E. coli*, and *Citrobacter* and has been evolutionarily maintained, indicating an important functional role.

Although SeaB is a clear positional orthologue of *E. coli* EhaB, our data reveal functional differences between the two proteins. Like EhaB, SeaB expressed in *E. coli* promotes biofilm formation and mediates bacterial aggregation; however, it displays distinct adhesion properties. We detected binding of SeaB to ECM components, similar to EhaB, but with phenotypic differences; SeaB preferentially binds to collagen III and fibronectin, whereas EhaB preferentially bound collagen I and laminin [26]. These differences in binding affinities likely stem from divergence in the passenger domain, which only share 49.7% identity, whereas the translocator and signal peptide sequences are more conserved (93.8% and 92.5% identity, respectively). The adhesion and biofilm-forming phenotypes are only observed in the *E. coli* BL21 strains overexpressing SeaB and EhaB, but not in their respective parent strains. The absence of a phenotype could be due to the presence of bacterial surface structures. Other studies have shown that the presence of surface structures such as fimbriae, O-antigen, lipopolysaccharide (LPS) and capsule abrogated the autoaggregation phenotype of T5SS antigens such as Ag 43, SadA and EhaA [24, 35, 48]. ECM binding antigens have also been associated with adhesion to and invasion of host cells. Although no specific interaction with CaCo2 cells was detected in this study SeaB may contribute to adhesion to other host cell types, as *Salmonella* exhibits cell type-dependent interactions during infection. Alternatively, its adherence function may be obscured by redundancy among other adhesins, and demonstrates the importance of redundancy in the need to bind extracellular matrix proteins [49].

ECM binding has been previously associated with bacterial virulence across multiple species [22, 50–53], implicating this function in host colonization and persistence. Moreover, biofilm development is a well-established factor in the chronic carriage and persistence of *S.* enterica [54, 55], and it is plausible that SeaB plays a role in this context. The demonstration that mutants in the periplasmic chaperones required for AT synthesis are also severely attenuated during murine infections indicates that ATs might be important for the pathogenesis of *Salmonella* [14]. Indeed, previous work by Chaudhuri *et al.* employing TraDIS revealed that disruption of *seaB* attenuates virulence in orally infected chickens, pigs, and cattle, but not in intravenously infected mice. The SalCom transcriptomic dataset from the Hinton lab previously identified SeaB as being upregulated under anaerobic shock [37]. Consistent with this our data further support a role for SeaB in infection in the gastrointestinal tract where conditions are hypoxic, potentially through mechanisms involving ECM binding and biofilm-mediated persistence. Interestingly, the absence of severe attenuation in the *seaB* mutant may reflect functional redundancy among *Salmonella* AT proteins or an a yet unestablished role for SeaB. Indeed, deletion of individual AT genes such as *sadA*, *misL*, or *shdA* has not resulted in major virulence defects [22–24], raising the possibility of overlapping or compensatory roles. However, loss of several of these proteins does result in diminished faecal shedding [23], which is an important facet in transmission of *Salmonella* to new hosts. Our findings are consistent with this hypothesis, suggesting that the full contribution of SeaB to pathogenesis may only be apparent when multiple ATs are simultaneously disrupted. Ongoing studies in our laboratory are aimed at dissecting this redundancy by generating multi-AT mutants.

In addition to its potential role in virulence, we investigated the immunogenicity of SeaB as a vaccine candidate. Structural modelling revealed similarity between SeaB and Pertactin, a key antigen in the acellular pertussis vaccine [19]. Immunization with alum-precipitated recombinant SeaB elicited robust IgG and IgM responses and conferred protection against *Salmonella* challenge in mice. These findings position SeaB as a promising component for inclusion in future *Salmonella* subunit vaccines. Importantly, our prior studies demonstrated that murine complement exhibits limited bactericidal activity against *Salmonella* [56], suggesting that antibody-mediated protection in this model may underestimate potential efficacy in humans. Nevertheless, the lack of a licensed protein-based monovalent vaccine against any Gram-negative bacterium underscores the challenge of developing effective subunit vaccines for these pathogens. The success of current vaccines, such as the acellular pertussis and Bexsero meningococcal vaccines, which employ multiple antigens, supports a multicomponent strategy for *Salmonella* as well. The specific immune pathways activated by SeaB and other T5SS proteins remain to be fully elucidated. Structural homologues of SeaB, including Pertactin and Filamentous Hemagglutinin, have been shown to modulate host immune responses by dampening inflammation, raising the possibility that SeaB may similarly influence the immune milieu during infection [57]. Furthermore, other *Salmonella* antigens have been shown to interact with distinct host cell types in diverse ways, thereby modulating the quality and efficacy of the antibody-mediated protective response [58, 59]. Identifying the innate and adaptive immune components engaged by these antigens will be critical for optimizing vaccine formulations.

In conclusion, our findings support a role for SeaB in biofilm formation, ECM binding, and virulence during oral infection, and demonstrate its potential as a protective antigen. However, functional redundancy among ATs and the complexity of the immune response to *Salmonella* suggest that a successful subunit vaccine will likely require targeting multiple antigens. Further studies exploring the combinatorial effects of AT deletions and multivalent vaccine approaches are warranted.

## ACKNOWLEDGMENTS

We acknowledge the support of the University of Birmingham MRC Scholarship to Faye Morris and a University of Queensland scholarship to Rochelle Da Costa. We also acknowledge the Australian Research Council Discovery Project for support of Jessica Rooke. We acknowledge Dhaarini Ragunathan for help in the construction of mutants.

---

**Figure S1.**
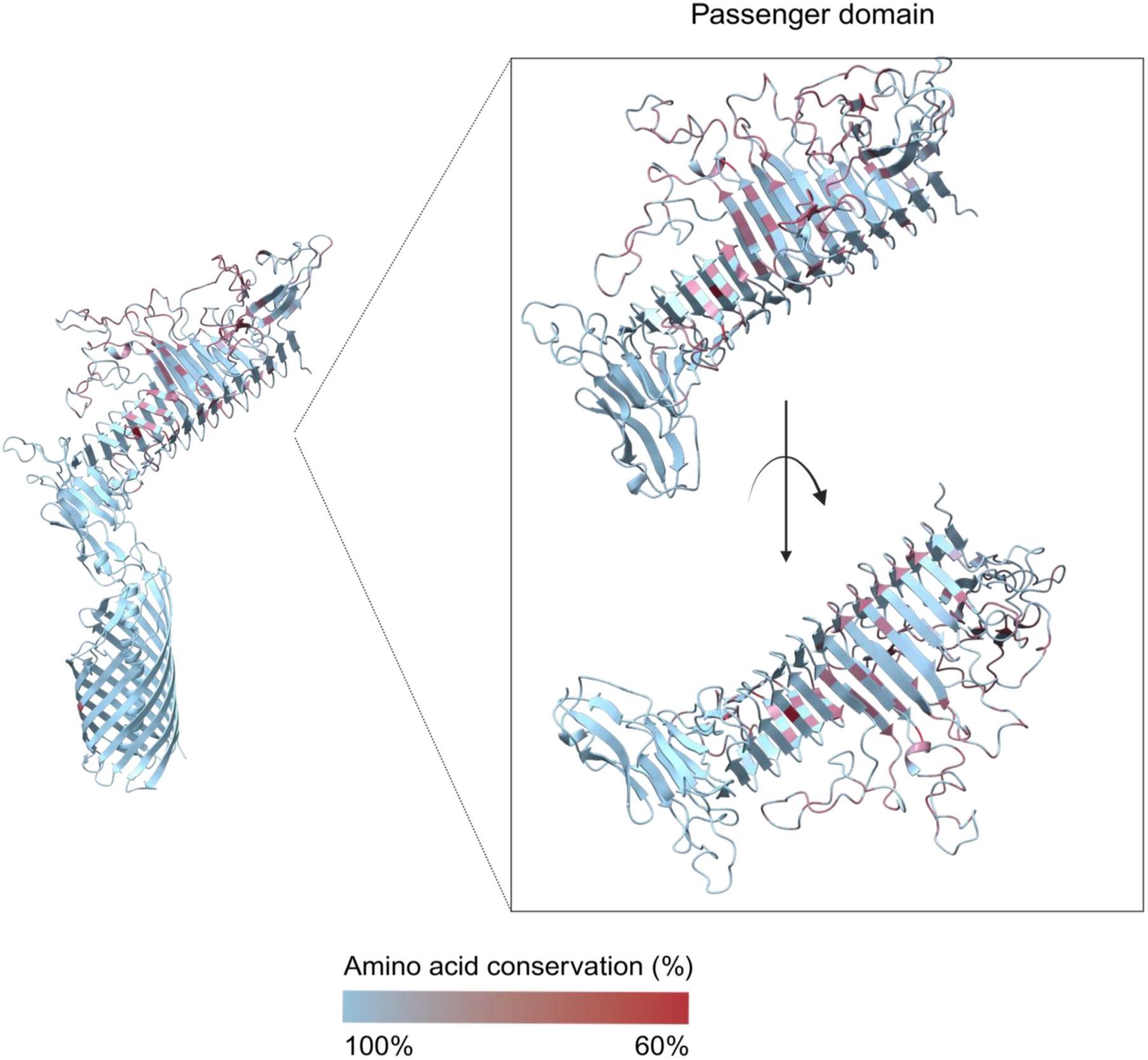
Structure of SeaB. Alphafold structure of SeaB with highly conserved regions shown in blue. Amino acid conservation for each residue was calculated using exact identity with the R package bio3d (v2.4-5) [60], and percent conservation between 60% to 100% was mapped onto the predicted structure of SeaB using ChimeraX (v1.10), with alignment gaps excluded from visualisation. The predicted domain structure of SeaB has a passenger domain similar to the solved crystal structure of Pertactin with a right-handed β helix, an autochaperone domain, and a 12 stranded C-terminal β-barrel.

**Figure S2.**
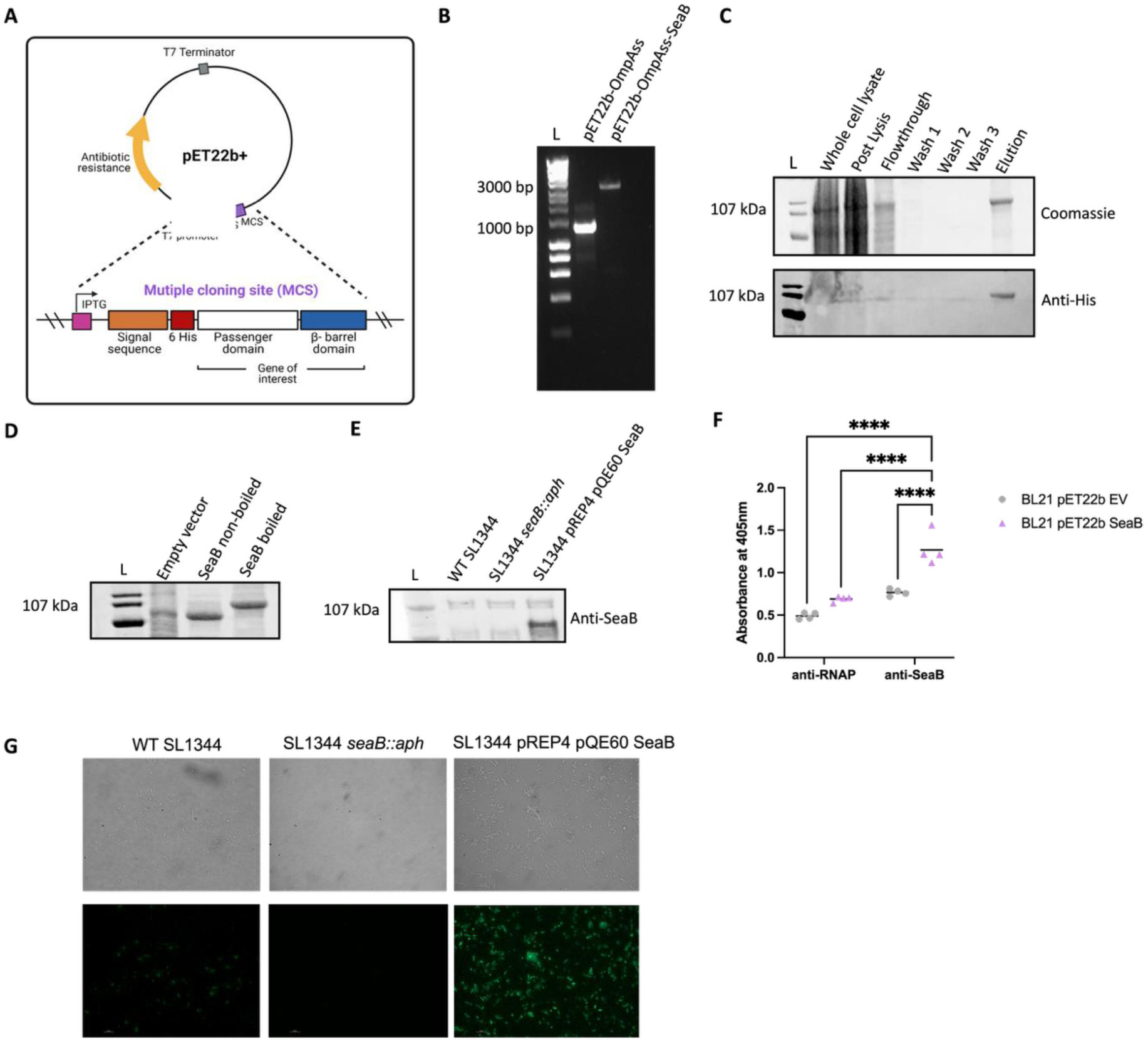
Cloning, purification, and localisation of SeaB. (A) Overview of the pET22b+ vector used to clone SeaB. OmpA signal sequence was first cloned into the commercially available overexpression vector pET22b+ between the *Nco*I and *Nde*I restriction sites. The passenger domain and β-barrel domain of SeaB was cloned into pET22b+ overexpression vector between *Not*I and *Kpn*I restriction sites in frame with OmpA signal sequence. (B) Gel image of OmpA ss cloned into pET22b+ (left) and SeaB cloned into pET22b+ (right). (C) *E. coli* BL21 P2 encoding SeaB was induced 50 µM IPTG for 16 h at 16°C. Samples were lysed using the Avestin C3. Following lysis, the protein was incubated with DDM overnight. SamplesSamples were purified by Nickel affinity chromatography and analysed by SDS-PAGE (above) and Western immunoblot using anti-His antibodies (below). (D) Purified protein samples (boiled and non-boiled) were run on an SDS-PAGE gel and visualised using Bio-Rad ChemiDoc MP Imaging system. (E) Expression of SeaB using anti-SeaB antibodies. (F) Surface localisation of purified SeaB. (G) Localization of SeaB on the cell surface. Strains were probed with anti-SeaB primary antibody followed by GAR Alexa Fluor 488. Statistical significance was determined using a two-way ANOVA Sidak’s multiple comparisons test (**** p<0.0001).

**Figure S3.**
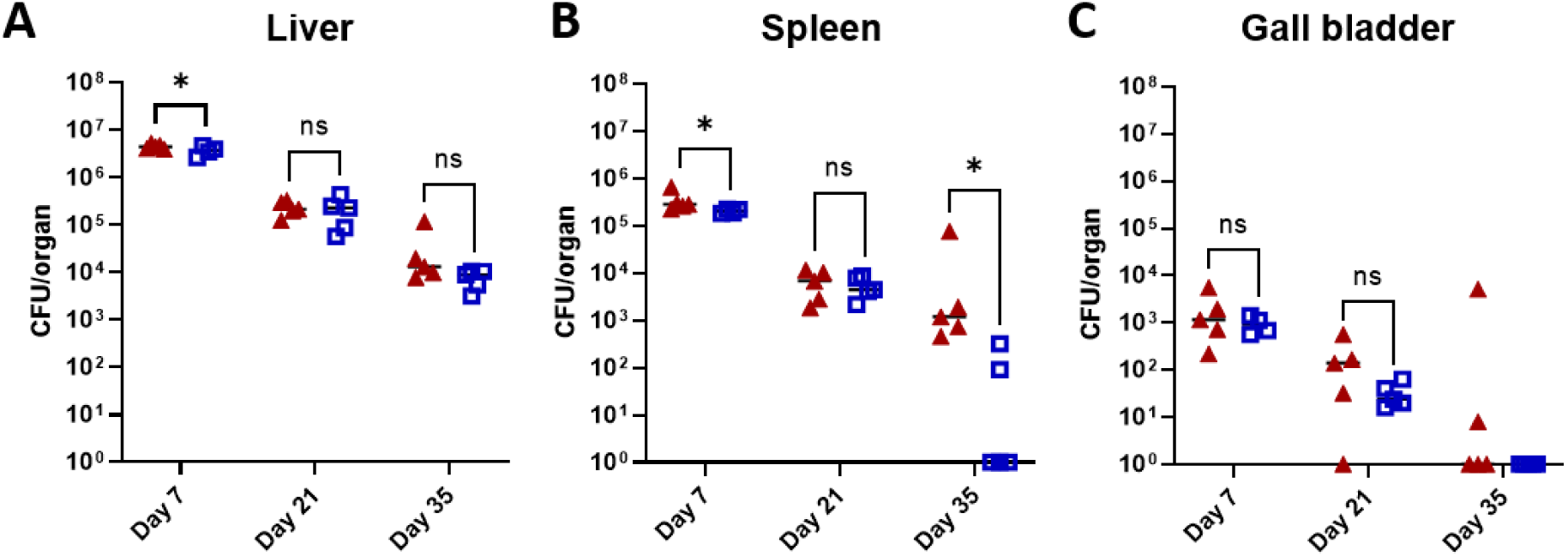
**Systemic infection with *Salmonella.*** Bacterial burdens in the (A) liver, (B) spleen and (C) gall bladder of mice infected for 35 days with WT SL3261 (red closed triangles) and SL3261 *seaB::aph* (blue open boxes). Mice were culled at days 7, 21 and 35 post infection. Statistical significance was determined using Mann-Whitney non-parametric test with correction for multiple comparisons (ns p>0.05, and *p<0.05).

**Table S1:**
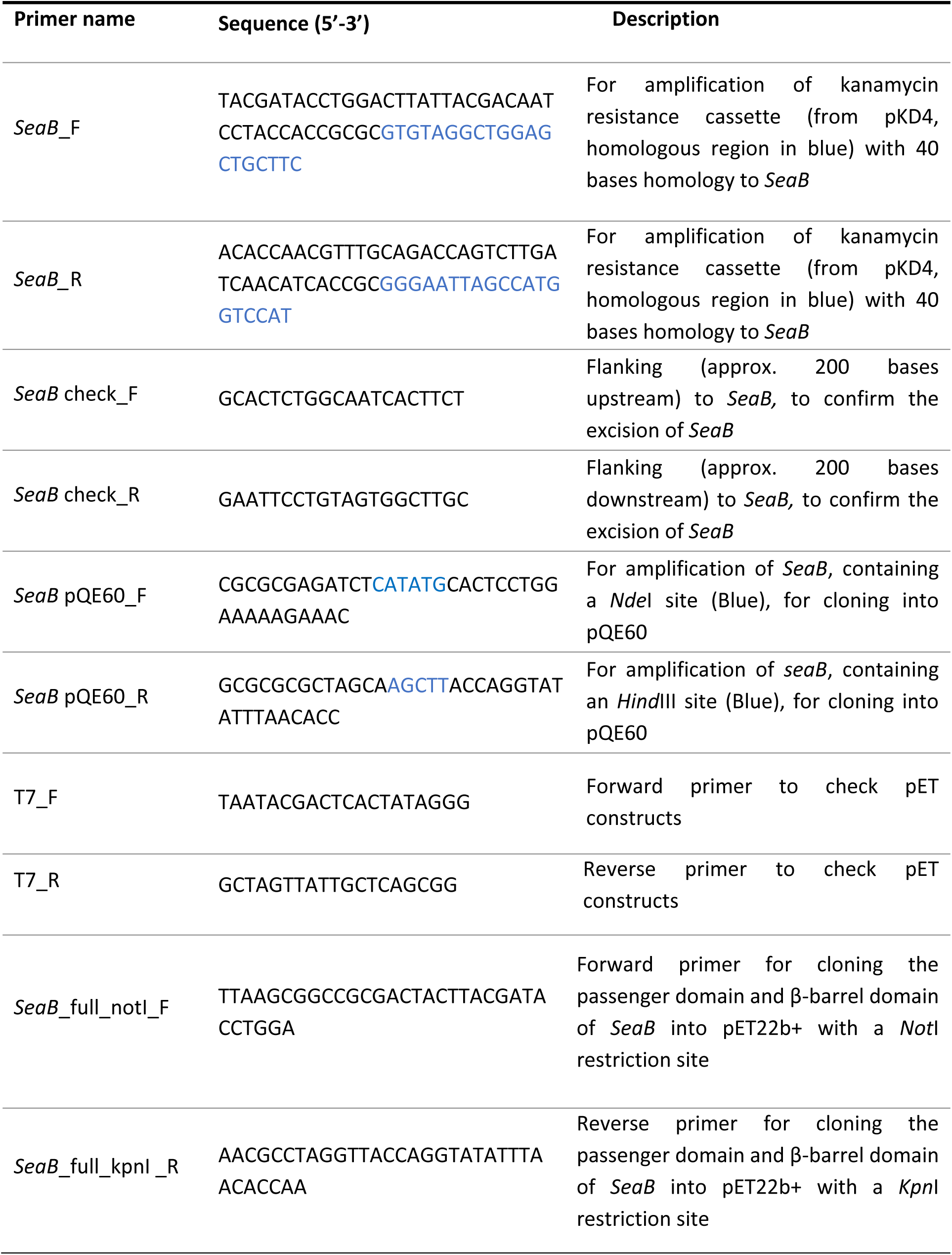
List of primers.

